# PlantCADB: A comprehensive plant chromatin accessibility database

**DOI:** 10.1101/2022.06.15.496248

**Authors:** Ke Ding, Shanwen Sun, Chaoyue Long, Yang Luo, Jingwen Zhai, Yixiao Zhai, Guohua Wang

**Affiliations:** State Key Laboratory of Tree Genetics and Breeding, Northeast Forestry University, Harbin 150040, China; College of Information and Computer Engineering, Northeast Forestry University, Harbin 150040, China; College of Life Science, Northeast Forestry University, Harbin 150040, China

**Author notes:** Corresponding author. (Guohua Wang).

**Keywords:** chromatin accessibility, plant, transcription factors footprint, regulatory network, stress response

## Abstract

Chromatin accessibility landscapes are essential for detecting regulatory elements, illustrating the corresponding regulatory networks, and, ultimately, understanding the molecular bases underlying key biological processes. With the advancement of sequencing technologies, a large volume of chromatin accessibility data has been accumulated and integrated in humans and other mammals. These data have greatly advanced the study of disease pathogenesis, cancer survival prognosis, and tissue development. To advance the understanding of molecular mechanisms regulating plant key traits and biological processes, we developed a comprehensive plant chromatin accessibility database (PlantCADB, https://bioinfor.nefu.edu.cn/PlantCADB/) from 649 samples of 37 species. Among these samples, 159 are abiotic stress-related (including heat, cold, drought, salt, etc.), 232 are development-related and 376 are tissue-specific. Overall, 18,339,426 accessible chromatin regions (ACRs) were compiled. These ACRs were annotated with genomic information, associated genes, transcription factors footprint, motif, and SNPs. Additionally, PlantCADB provides various tools to visualize ACRs and corresponding annotations. It thus forms an integrated, annotated, and analyzed plant-related chromatin accessibility information which can aid to better understand genetic regulatory networks underlying development, important traits, stress adaptions, and evolution.

## Introduction

Eukaryotic chromatin is an un-uniformly compacted complex of DNA and proteins. Its physical compactness is referred as chromatin accessibility, which is determined by nucleosome occupancy, topological structure, post-translational chemical modifications and other chromatin binding factors. Less condensed chromatin forms an ACR on the genome that can be contacted by nuclear macromolecules. Previous study showed that although these regions account for only 2-3% of the total DNA sequence in human, they accommodate important *cis*-regulatory elements that capture about 94% all ENCODE transcription factors binding sites [1]. Mapping chromatin open landscape on a genome-wide scale is thus vital for detecting *cis*-regulatory elements and understanding the regulation of important biological processes [2, 3]. For example, Pajoro *et al*. constructed the dynamic regulatory networks during *Arabidopsis* flower development by monitoring the changes of chromatin accessibility and gene expressions [4]. This work helps to illustrate mechanisms of MADS-domain transcription factors, the well-known master regulators, to regulate development and organ specification [4]. Moreover, by mapping the ACRs of control and cold-stressed samples from three grasses, Han *et al*. found a significant enrichment of cold-induced ACRs adjacent to cold responsive genes and a high conservation of transcription factor (TF) binding motifs embedded in these regions, suggesting that common transcription factors may regulate the transcriptional adaptation to cold stresses across species [5]. In *Arabidopsis*, the phosphorylation of DEK3, a chromatin architectural protein, was found to alter nucleosome occupancy and chromatin accessibility and lead to changes in gene expression, ultimately promoting salt stress tolerance [6]. Together, these results illustrate the importance of chromatin accessibility data in addressing key issues related to the molecular regulation of biological processes, development and stress adaptations.

Experimental methods to identify such chromatin accessibility regions throughout the genome rely on combining enzymatic digestion of nuclear DNA and high-throughput sequencing, including DNase I Hypersensitivity sequencing (DNase-seq) [7, 8], Microccocal nuclease sequencing (MNase-seq) [9], Assay for Targeting Accessible-Chromatin with sequencing (ATAC-seq) [10, 11], and Formaldehyde-assisted isolation of regulatory elements sequencing (FAIRE-seq) [12]. The main idea of DNase-seq and MNase-seq is to use enzymes to cut DNA double strands and the sequencing result is accessible regions of chromatin. They are widely used in the analysis of cell-specific chromatin accessibility and to investigate the relationship between chromatin accessibility and gene expression [8, 9]. ATAC-seq detects the regions bound by transcription factors or occupied by nucleosomes. It is faster and more sensitive than DNase-seq and MNase-seq [13, 14]. FAIRE-seq overcomes the enzymatic cleavage preference that may present in the above methods and directly detects DNA sequences occupied by nucleosomes [12]. In general, these methods can accurately and sensitively reflect the open landscape of chromatin.

With declining sequencing costs and the development of easy-to-use library construction tools, research on chromatin accessibility in humans, other mammals, and plants have become very mature. To fully exploit the large volume of chromatin accessibility data, several databases have been offered to the public. For instance, the Cistrome Data Browser compiled chromatin accessibility data in human and mouse with homeopathic regulatory information [15]. It helped to illustrate the regulatory relationship and survival prognosis in human cancer [16, 17]. The online database ‘Brain Open Chromatin Atlas (BOCA)’ provides an accessible atlas of human brain chromatin [18]. This database has greatly advanced research on Alzheimer’s disease, neuropsychiatric disorders and human brain development [19-22]. Chen *et al*. further developed OpenAnnotate to assess the chromatin accessibility of large-scale genomic regions based on features extracted from public chromatin accessibility data [23]. It builds a more comprehensive perspective to understand regulatory mechanisms in humans and mouse [23]. These cases demonstrate the power of compiled chromatin accessibility database to facilitate the understanding of gene regulation in mammals and accelerate the study of pathogenesis in diseases.

Deeper analyses of chromatin accessibility data are of paramount importance for plants as well. For example, Tannenbaum *et al*. found that performing motif enrichment analysis on root-specific accessible regions enabled them to discover root-specific TFs that are related to root development [24]. By integrating TF CHIP-seq and ATAC-seq data, Tu *et al*. found that TF co-binding could be key for transcriptional regulation of *Zea mays*, which was conducive to the rapid diversification of the regulatory network during speciation [25]. Moreover, chromatin accessibility is crucial to illustrating how genetic variation, such as SNPs in non-coding regions, leads to plants functions, adaptation and ultimately evolution. However, public databases on plant chromatin accessibility and its subsequent analyses are missing. To provide a comprehensive chromatin accessibility data analysis platform for plants, we developed PlantCADB. It compiled a large number of available open chromatin landscape resources and annotated their potential roles in regulation. In total, 18,339,426 ACRs from more than 600 samples of 37 species are available in the database (Table 1). Among these samples, 159 are abiotic stress-related (including heat, cold, drought, salt, etc.), 232 are development-related and 376 are tissue-specific. ACRs genome annotation, associated gene annotation, SNP annotation, TF footprint analysis and motif scanning analysis are offered. Users can also perform data search, region visualization, ACRs difference analysis and overlap analysis on the web page. It additionally provides data quality control analysis, statistics and download functions. These characteristics form an integrated, annotated and analyzed chromatin accessibility information which can aid to better understand the molecular mechanisms underlying key traits, biological processes, development and stress adaptations in plants.

**Table 1.** Chromatin accessibility data summary.

| Class | Species | ATAC-seq |  | DNase-seq |  | FAIRE-seq | MNase-seq |  | Total |
| --- | --- | --- | --- | --- | --- | --- | --- | --- | --- |
|  |  | GEO | SRA | GEO | SRA | GEO | GEO | SRA |  |
| **Dicotyledoneae** | *Arabidopsis thaliana* | 85 | 27 | 48 | 12 | 15 | 3 |  | 190 |
|  | *Solanum lycopersicum* | 16 |  |  | 77 |  |  |  | 93 |
|  | *Cucumis melo* |  |  |  | 30 |  |  |  | 30 |
|  | *Prunus persica* |  |  |  | 21 |  |  |  | 21 |
|  | *Populus trichocarpa* | 18 |  |  |  |  |  |  | 18 |
|  | *Pyrus x bretschneideri* |  |  |  | 14 |  |  |  | 14 |
|  | *Citrullus lanatus* |  |  |  | 10 |  |  |  | 10 |
|  | *Cucumis sativus* |  |  |  | 10 |  |  |  | 10 |
|  | *Medicago truncatula* | 9 |  |  |  |  |  |  | 9 |
|  | *Solanum pennellii* | 8 |  |  |  |  |  |  | 8 |
|  | *Vitis vinifera* |  |  |  | 8 |  |  |  | 8 |
|  | *Glycine max* | 7 |  |  |  |  |  |  | 7 |
|  | *Malus domestica* |  |  |  | 7 |  |  |  | 7 |
|  | *Fragaria vesca* |  |  |  | 7 |  |  |  | 7 |
|  | *Solanum phureja* |  |  |  | 6 |  |  |  | 6 |
|  | *Carica papaya* |  |  |  | 6 |  |  |  | 6 |
|  | *Arachis hypogaea* |  | 4 |  |  |  |  |  | 4 |
|  | *Eutrema salsugineum* | 3 |  |  |  |  |  |  | 3 |
|  | *Phaseolus vulgaris* | 3 |  |  |  |  |  |  | 3 |
|  | *Gossypium arboreum* |  |  |  | 1 |  |  |  | 1 |
|  | *Gossypium barbadense* |  |  |  | 1 |  |  |  | 1 |
|  | *Gossypium hirsutum* |  |  |  |  |  |  | 1 | 1 |
|  | *Gossypium raimondii* |  |  |  | 1 |  |  |  | 1 |
|  | *Eucalyptus grandis* |  |  |  | 12 |  |  |  | 12 |
| **Monocotyledoneae** | *Zea mays* | 23 | 13 | 11 | 2 |  |  |  | 49 |
|  | *Oryza sativa* | 58 | 2 |  |  |  |  |  | 60 |
|  | *Sorghum bicolor* | 3 | 3 | 6 |  |  |  |  | 12 |
|  | *Ananas comosus* |  |  |  | 12 |  |  |  | 12 |
|  | *Setaria italica* |  | 2 | 6 |  |  |  |  | 8 |
|  | *Brachypodium distachyon* | 3 |  | 3 |  |  |  |  | 6 |
|  | *Musa acuminata* |  |  |  | 6 |  |  |  | 6 |
|  | *Hordeum vulgare* | 3 |  |  |  |  |  |  | 3 |
|  | *Setaria viridis* | 3 |  |  |  |  |  |  | 3 |
|  | *Asparagus officinalis* | 3 |  |  |  |  |  |  | 3 |
|  | *Spirodela polyrhiza* | 3 |  |  |  |  |  |  | 3 |
|  | *Triticum aestivum* | 1 |  |  |  |  |  |  | 1 |
| **Hepaticae** | *Marchantia polymorpha* |  | 2 |  |  |  |  |  | 2 |
| **Total** |  | **249** | **53** | **74** | **243** | **15** | **3** | **1** | **638** |

## Result

### Statistics of PlantCADB

In the current version of PlantCADB, we collected a total of 649 samples from 37 species (Figure 1A, see Table 1 for details), covering four types of sequencing data from 19 tissues (Figure 1B). The plant species include angiosperms (monocotyledons and dicotyledons) and bryophytes with different genome sizes (122 Mb-14790 Mb), genome structures and gene densities (7.5-124 genes/Mb). Overall, 18,339,426 ACRs were identified (7,972,702 from ATAC-seq; 9,472,599 from DNase-seq; 568,255 from FAIRE-seq; 325,870 from MNase-seq). The total sequence length of ACRs in each species ranges from 0.4 to 104.3 Mb and accounts for 0.003-11.3% of the genome size across species. The numbers of ACRs, its total sequence length and the percentage of the genome size occupied by all ACRs in each species do not significantly increase with the increased genome size (Spearman rank correlation ρ_number_ = 0.19, ρ_length_ = 0.14, ρ_percent_ = -0.48, all p-values > 0.05; Supplement Figure 1). After motif scanning analysis on 624 samples of 33 species (four species without available motif data were excluded from the analysis), we found that approximately 99.1% of ACRs have the potential to be bound by TFs. After analyzing the distribution of SNPs in different sequencing data types of each species from a total of eight species for which SNP data is available, we found that the density of SNPs in ACRs is significantly higher than whole genome (the ratios between densities ranging from 1.43 to 25.82; Figure 1C).

**Figure 1.**
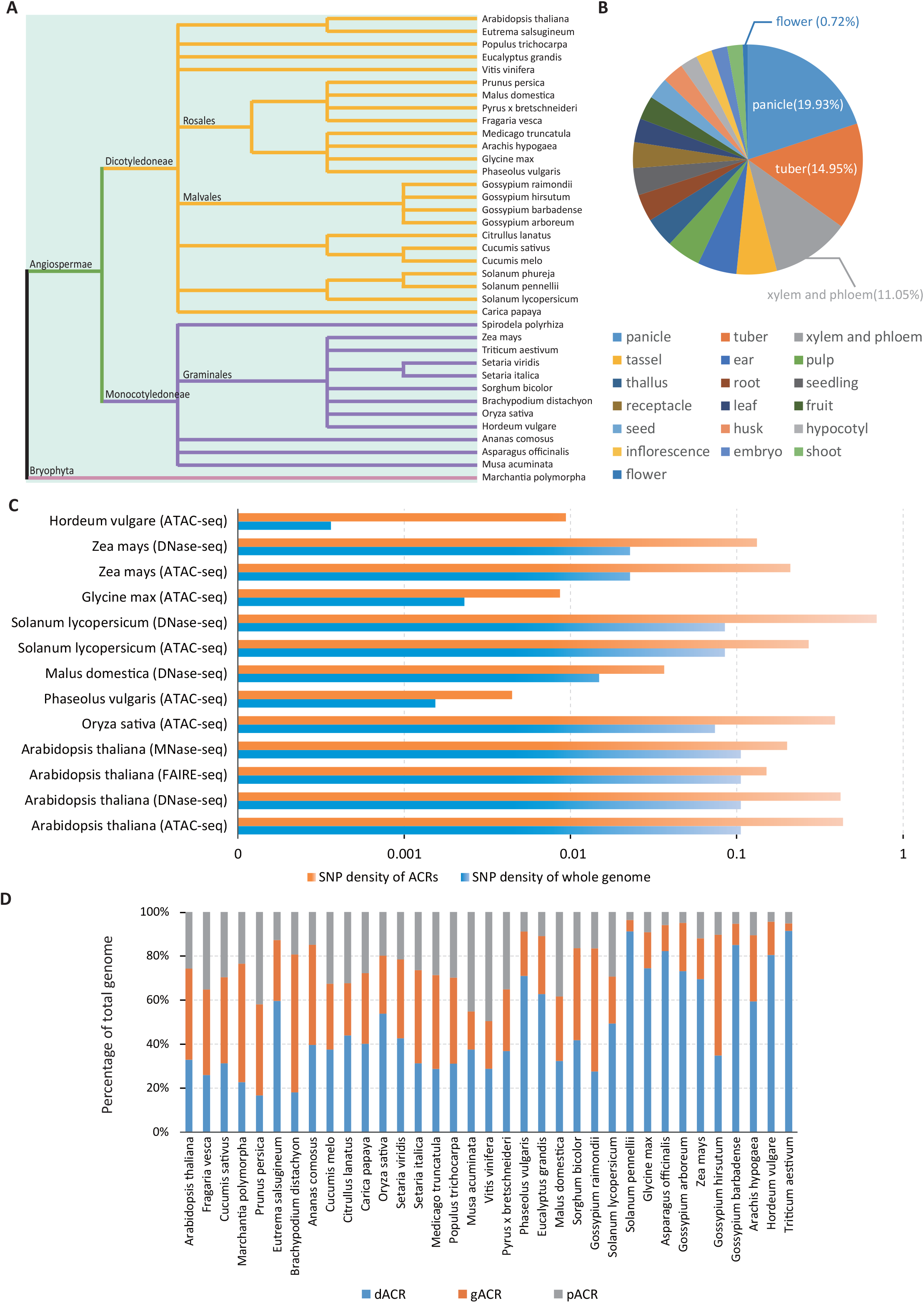
Characteristics of PlantCADB and ACRs. **A**. A phylogenetic map of the plant species that were investigated. **B**. A pie chart of the average number of ACRs in different tissues. **C**. The density of SNPs in eight species in the whole genome and in ACRs. **D**. The average percentage of dACRs, gACRs and pACRs in the genome of each species (species are ordered based on the reference-genome size).

ACRs were further classified into genic (gACRs, overlapping with gene), proximal (pACRs, within 2kb from a gene) and distal (dACRs, distance from a gene > 2kb) according to distance between ACRs and the nearest gene [26]. We found that the number of dACRs (8,013,939) is positively correlated with increasing genome sizes (Spearman rank correlation ρ = 0.66, p-value < 0.001; Figure 1D), while the numbers of gACRs (5,488,948) and pACRs (4,157,698) are both negatively correlated with increasing genome sizes (Spearman rank correlation ρ_pACRs_ = -0.62, ρ_gACRs_ = -0.57, both p-values < 0.001; Figure 1D). In addition, the numbers of the three ACRs types are also significantly correlated with gene density across species (Spearman rank correlation ρ_dACRs_ = -0.73, ρ_pACRs_ = 0.63, ρ_gACRs_ = 0.62, all p-values < 0.001; Supplement Figure 2).

**Figure 2.**
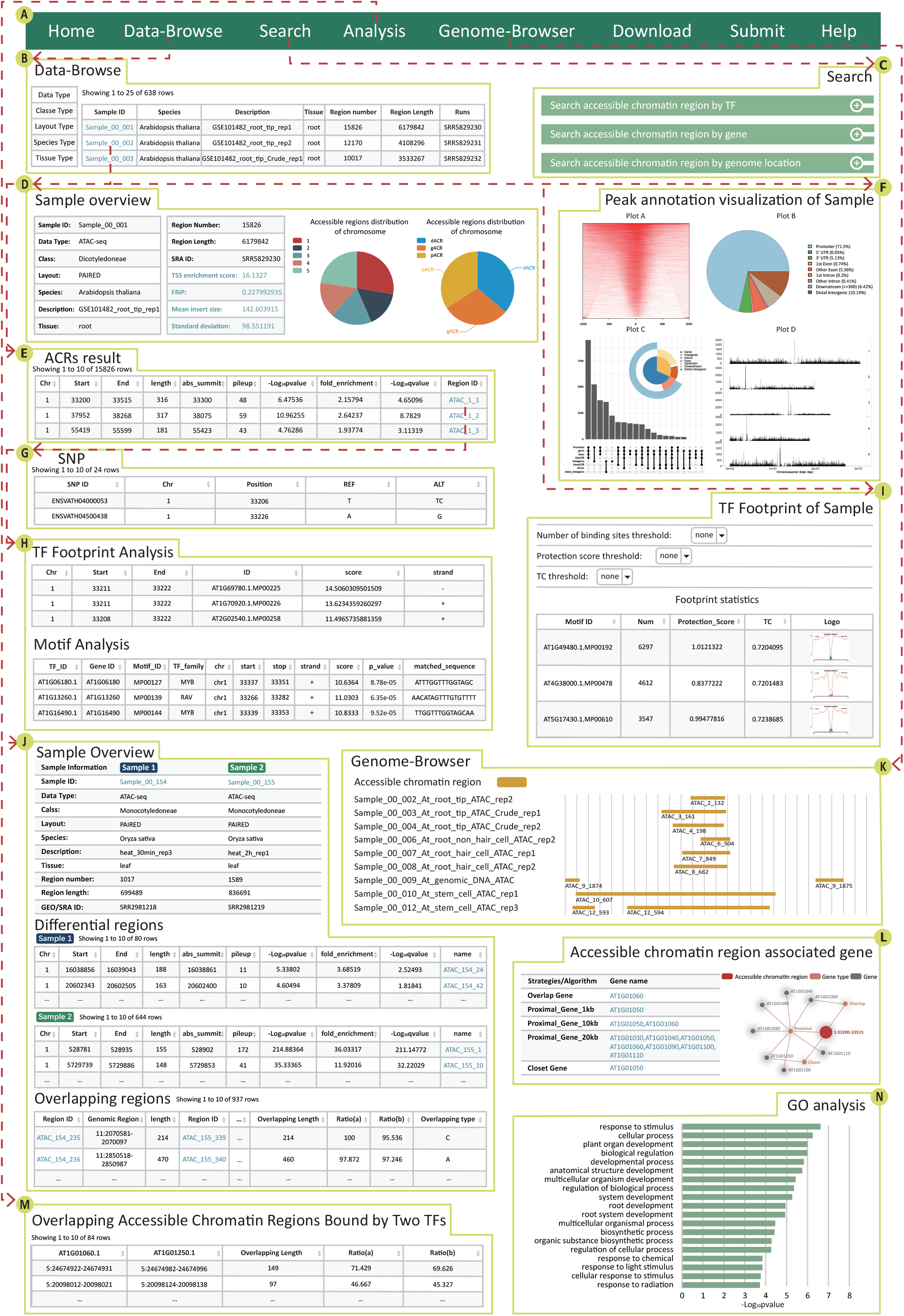
The main functions and usage of PlantCADB. **A**. The navigation bar in PlantCADB. **B**. Browsing the sample details. **C**. Users can query chromatin accessibility regions through three paths: ‘Search by TF’, ‘Search by gene’ and ‘Search by genomic location’. **D**. Sample information including sample ID, data type, class, layout, species, description, tissue, region number, region length, SRA ID, quality control report and pie chart of ACRs distribution. **E**. Table of ACRs results for one sample including chromosome, start, end, length, abs_summit, pileup, -log_10_pvalue, fold_enrichment, -log_10_qvalue and region ID. **F**. Peak annotation visualization of a sample. **G**. The detailed information of SNP. **H**. The detailed information on each region of TF analysis including TF footprint and motif scanning. **I**. TF footprint statistics of sample. **J**. Analysis of differential and overlapping ACRs between two samples in same species. **K**. Visualization of genome browser. **L**. Potential ACRs associated genes that are identified by three strategies. Their relationships are displayed using a network diagram. **M**. Analysis of overlapping ACRs bound by two TFs in same species. **N**. Gene ontology analysis of biological process associated with ACRs of MYC2 transcription factor in *Arabidopsis thaliana*.

### Web interface of PlantCADB

#### A search interface for retrieving chromatin accessibility regions

PlantCADB offers three user-friendly search options to retrieve chromatin accessibility data (Figure 2C). With TF-based query (‘Search accessible chromatin region by TF’), users can obtain all ACRs that are potentially bound by the query TF by selecting species and TF ID. With gene-based query (‘Search accessible chromatin region by gene’), users can obtain all ACRs associated with the gene of interests after selecting species and identification strategy (OVERLAP, PROXIMAL and CLOSEST). With genomic region-based query (‘Search accessible chromatin region by genome location’), users can get all ACRs that overlap (at least 1kb) with the submitted region by selecting species and sample ID, and inputting genomic location.

#### A browsing interface for retrieving chromatin accessibility regions

Users can browse all ACRs belonging to a specific data type, taxon, layout (the construction method of sequencing library), species or tissue (Figure 2B). The result shows samples that match the filter conditions. All samples from four sequencing technologies were named with different prefixes (‘Sample_00’ for ATAC-seq, ‘Sample_01’ for DNase-seq, ‘Sample_02’ for FAIRE-seq and ‘Sample_03’ for MNase-seq). User can further click the ‘Sample ID’ to get the detailed information of this sample, including sample overview, ACRs result table, TF footprint annotation and peak annotation visualization (Figure 2D, E, F and I). The sample overview provides general information, values of four quality control (QC) metrics, measurement indicators and pie charts of statistical information about ACRs of this sample (Figure 2D). Based on two statistical pie charts, users can view the distribution of ACRs on each chromosome and the number of various types of ACRs. ACRs result table shows all ACRs of the sample. It describes region ID, genome location, region length, summit site (abs_summit), summit height (pileup), fold enrichment and -log_10_pvalue (Figure 2E). The TF footprint annotation shows the results of HINT software analysis. Tag Count (TC), protection score, number of binding sites and footprint logo were identified for each sample. We also offer a ‘threshold’ option, which allows users to screen for TFs with high activity by setting thresholds (Figure 2I). The peak annotation visualization displays the annotation information of ACRs in different ways, including peak distribution heat maps near transcription start site (TSS) regions, pie charts annotating genomic features, panoramic maps of ACRs in the genome distribution and more detailed combination chart (Figure 2F).

To view more detailed information of the designated ACR, we provide TF footprint corresponding scores, the results of motif scanning, SNP and associated genes (Figure 2G, H and L). Motif scanning includes location information, sequence scores and matched sequences. The associated genes are further classified into overlap genes, proximal genes within ±1kb, proximal genes within ±10kb, proximal genes within ±20kb and the closest genes.

#### Online analysis tools

Analyzing the dynamics of chromatin accessibility can allow us to understand the changes of molecular regulation in response to developmental cues and external stimuli [27]. PlantCADB provides two online analysis tools. The first one is differential-overlapping analysis of ACRs. In the ‘Analysis of differential-overlapping accessible chromatin region’ panel, user can submit two ‘Sample ID’ of interest and get the analyzed differential and overlapping ACRs between the two samples (Figure 2J). For overlapping regions (at least 1bp overlap between them), we divide them into four types. Type A indicates that the right wing of ACR in ‘Sample 1’ overlaps at least 1bp with the left wing of ACR in ‘Sample 2’. Type B is opposite to type A, indicating that the left wing of ACR in ‘Sample 1’ overlaps the right wing of ACR in ‘Sample 2’. Type C means that a certain ACR of ‘Sample 1’ is completely covered by a certain ACR of ‘Sample 2’. Type D is completely opposite to type C, which means that a certain ACR of ‘Sample 1’ completely covers the ACR of ‘Sample 2’. We additionally provide the genomic positions of the two overlapping regions, overlap position, overlap length, and ratios of overlap (ratio of overlapping areas in each ACR). For differential regions, we define them as different regions and output these regions of the two samples respectively.

Users can also analyze the overlapping ACRs bound by two TFs. In the ‘Analysis of overlapping accessible chromatin regions bound by two TFs’ panel, users can submit two interested ‘TF name’ and ‘window length’ (Figure 2M). The tool can fetch all regions that are bound by both TFs and calculate the two overlapping areas according to the submitted window length. The results of analysis are briefly displayed in a table, including the genomic location of TF, length of the overlap region and overlap rates (ratio of the overlap length to total length, where the total length = the length of the TF bound to the genome location + 2 * the length of the window).

#### Data visualization and personalized genome browser

To help users better view the genomic information of chromatin accessibility, we also provide a personalized genome browser, which is developed using the latest version of JBrowser2 [28]. Users can intuitively see the positional relationship between chromatin accessibility and nearby genes, mRNA, tRNA, lncRNA and rRNA, and other genomic fragments (Figure 2K). In addition, we also provide annotated pie charts and distribution maps of ACRs, and the network between ACRs and genes drawn online using Echarts software.

### Case studies

To provide an example of how regulators can be used in PlantCADB to retrieve the putative corresponding regulatory network, the bHLH TF family transcription factor MYC2 (AT1G32640.1) in *Arabidopsis thaliana* is used as input to our database for ‘Search by TF’. After clicking the ‘Start search’ button, a total of 922 ACRs regions with 110 nearest neighbor genes that are potentially bound by MYC2 were retrieved. In order to characterize potential functions of these associated genes, we performed a gene ontology enrichment analysis using PANTHER. 19 significant biological processes were obtained (FDR P < 0.05, Figure 2N), including root development, response to light stimulus, organic substance biosynthetic process, response to chemical, cellular response to stimulus and regulation of cellular process. These results are consistent with previous findings which indicate that MYC2 is an important regulator of lateral root formation and light responses [29, 30] and suggest that MYC2 may perform other regulatory roles in biosynthesis or responses to stimulus.

Analyzing the changes of chromatin accessibility can help to reveal the dynamics of transcriptional regulatory landscape during the development or in response to external stimuli [31]. Here is an example to show how to use overlapping-differential analysis tools to identify dynamic changes in chromatin accessibility to respond to heat. In ‘Analysis of differential-overlapping accessible chromatin region’ interface, after selecting ‘*Oryza sativa*’ species, ‘abiotic stress’ experimental classify, ‘Sample_00_154:heat_30min_3’ and ‘Sample_00_155:heat_2h_1’, click the ‘Start analysis’ button to start analysis. These two samples are 14 days old rice leaves (second leaf) in heat stress (transferred from 30□ to 40□) for 0.5h and 2h, respectively. The upper interface shows the detailed information of the two samples as well as the pie chart of overlap rate (percentage of overlapping ACRs of all ACRs in this sample) and difference rate (percentage of difference ACRs of all ACRs in this sample). The lower interface displays differential regions and overlapping regions between the two samples, respectively. In differential regions part, ‘Sample_00_155:heat_2h_1’ has more ACRs than ‘Sample_00_154:heat_30min_3’. In the overlapping regions section, we divided overlapping types into four types. In this example, type C accounts for the most (∼51.5%) and type D the least (∼15.7%). Type A and type B account for ∼16.2% and ∼16.8% respectively. These results suggest that chromatin accessibility landscape expands with increasing exposure to high temperature in *Oryza sativa*, which cooccurs with the expression of about 500 more genes [32], indicating the transcriptional reprogramming in response to the heat stress.

To compare tissue-specific ACRs, we used ‘Sample_00_237:maize_B73Leaf_rep1’ (leaf tissue) and ‘Sample_00_238:maize_B73Ear_rep1’ (ear tissue) from *Zea mays*. We identified a total of 67,532 and 34,270 ACRs in leaf and ear, respectively. Among them, 23,443 are shared ACRs, and 43,949 and 9,707 are tissue-specific ACRs in leaf and ear, respectively. In order to characterize the potential functions of ACRs, we extracted the nearest neighbor gene of each ACRs for gene ontology enrichment analysis. For shared ACRs, the results of the Gene Ontology (GO) analysis included 87 terms, such as regulation of RNA biosynthetic process, regulation of nucleic acid-templated transcription, and regulation of transcription, DNA-templated. For tissue-specific ACRs, we obtained 19,450 and 7,384 nearest neighbor genes from leaf and ear tissue, respectively. To further ensure that these genes function in a tissue-specific manner, we integrated RNA-seq data from leaf and ear tissues. The corresponding RNA-seq [33] data was analyzed with limma test to obtain differentially expressed genes. 2,382 and 795 genes in the nearest neighbor genes are differentially expressed in leaf and ear, respectively. Interestingly, regulation of RNA biosynthetic process, regulation of nucleic acid-templated transcription and regulation of transcription, DNA-templated are the top three significant GO groups in both tissues, which suggest that different sets of ACRs and genes may play a significant role in regulating transcription in different tissues. Moreover, we found that the genes related to leaf-specific ACRs are heavily enriched in chloroplast rRNA processing and protein localization to chloroplast.

## Discussion

Profiling chromatin accessibility on a genome wide scale is widely used to understand the transcriptional regulation, tissue specificity, stress responses and developmental dynamics of plant [34]. For example, Wu *et al*. systematically studied the combined effects of multiple epigenome features on gene expression in *Arabidopsis thaliana* and *Oryza sativa* based on histone modifications and chromatin accessibility data [35]. Wang *et al*. studied ATAC-seq data at different stages of somatic embryogenesis induced by auxin and proved that auxin can rapidly reconnect the totipotent network of cells by altering chromatin accessibility in *Arabidopsis thaliana* [34]. Moreover, based on the dynamic analysis of chromatin accessibility, they also revealed the hierarchical gene regulatory network in the process of somatic embryogenesis [34]. These applications benefit from high genomic resolution of chromatin accessibility analysis, reasonable cost and the ability to process many samples in a fast manner. Database such as ENCODE [36, 37], TCGA [38] and Cistrome [15] all focus on providing original chromatin accessibility data in human and are being extensively used by tools to annotate cis-regulatory elements such as enhancers [39] and silencers [40]. There is currently no database that provides a collection of complete chromatin accessibility regions and detailed annotation information and analyses of ACRs in plants. Here, we provide the PlantCADB, which can make it easier for users to use ACRs and understand the underlying biological functions.

PlantCADB is a comprehensive database of ACRs, providing a convenient interface to browse, query, analysis, visualize and download ACRs. The advantages of PlantCADB include: (I) comprehensive ACRs of plant species; (II) inferred TF binding in ACRs using TF footprint analysis; (III) options to query the associated ACRs with user submitted genome location, gene or transcription factor; (IV) useful online analysis tools for ACRs such as ‘Analysis of differential-overlapping ACRs’ and ‘Analysis of overlapping ACRs bound by two TFs’; (V) personalized genome browser for intuitively viewing information of ACRs and adding other useful tracks; (VI) conveniently displaying and downloading of ACRs and related annotation information via interactive tables. As illustrated in three case studies, PlantCADB provides convenient tools to explore the relationship between genes, transcription factors and chromatin accessibility regions to decipher the key questions in plant science. Although till now plants chromatin accessibility are only assessed with bulk sequencing data, with the development of single-cell sequencing technology, users will soon be able to construct the epigenetic landscape of single cell and cell differentiation trajectories with tools such as scDEC [41], RA3 [42] and epiAnno [43]. PlantCADB will be updated in time to add new datasets and be applied to more plant species.

## Materials and methods

### Data collection and identification and classification of chromatin accessibility regions

To make full use of publicly available large-scale sequencing data, we manually collected all ATAC-seq, DNase-seq, MNase-seq and FAIRE-seq data related to plant from NCBI GEO/SRA [44]. These sequences are from 37 species and 19 tissues (Table 1). The reference genomes and corresponding genes of each species were downloaded from NCBI Genome (https://www.ncbi.nlm.nih.gov/genome/) and Ensembl Plants [45, 46].

The downloaded raw sequencing data were first converted into FASTQ format files using fastq-dump in NCBI SRA Toolkit (version 2.9.2; Figure 3). FastQC [47] was used to examine sequence quality, GC content, sequence length distribution, sequence duplication levels, overrepresentation sequences and contamination of adapters in the raw sequencing data for pre-alignment QC. Adapters and low-quality reads were removed using Trim Galore (version 0.6.6; Figure 3) with the following parameters ‘-q 20 --phred33 --stringency 3 --length 20 -e 0.1’. After that, sequences were mapped to the reference genome (Supplement Table 1) using Bowtie2 (version 2.4.2; Figure 3) [48] with default parameters.

**Figure 3.**
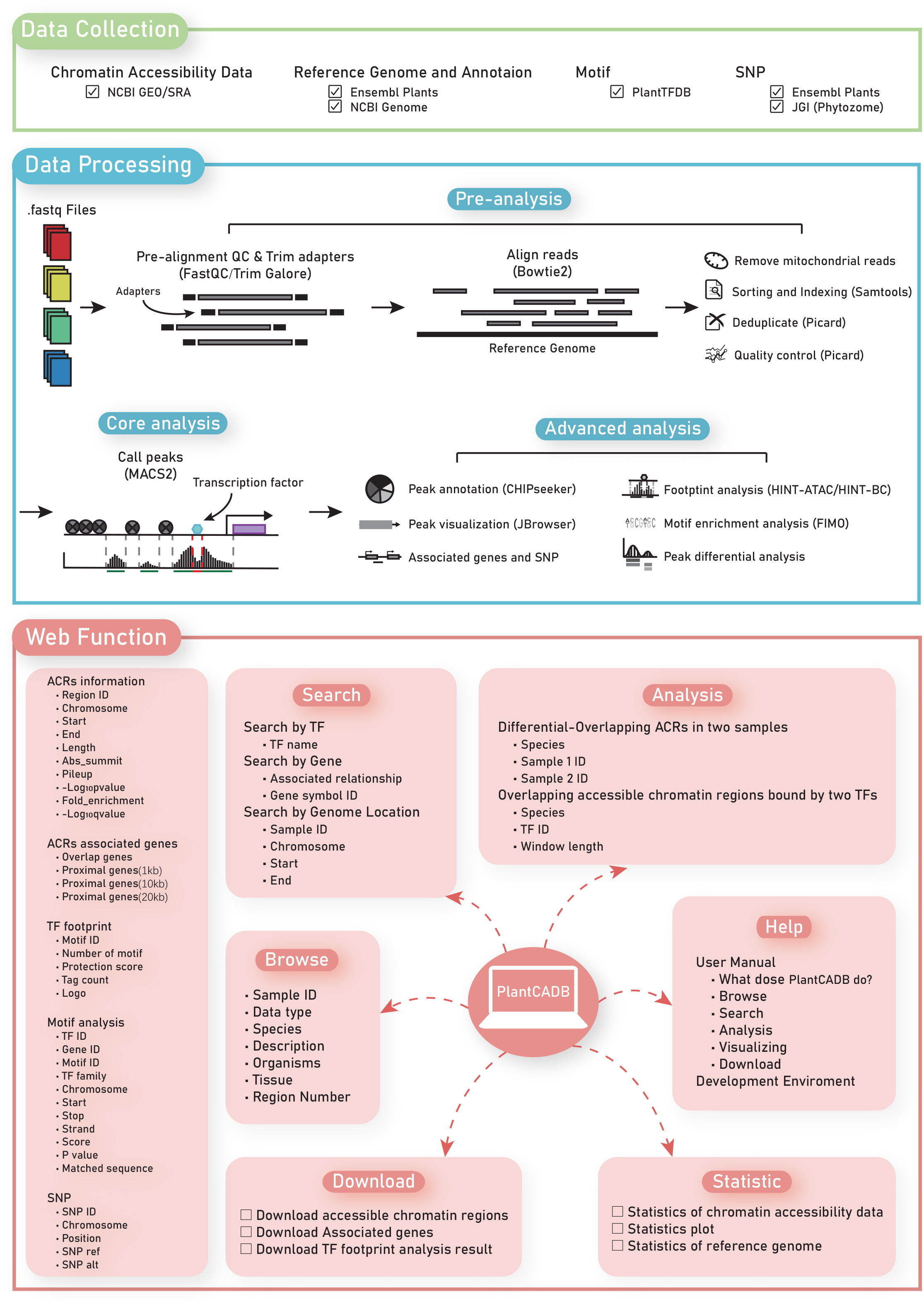
Database content and construction. PlantCADB calculated ACRs using ATAC-seq, DNase-seq, FAIRE-seq and MNase-seq data. Genetics annotations were collected or calculated including SNPs, TFBSs, TF footprint and associated genes. Users can query ACRs using three strategies: TF-based query, gene-based query and genome-location-based query. PlantCADB also includes online analyzing tools and personalized genome browser to discover potential biological effects of ACRs.

To improve the power of open chromatin detection and produce fewer false positives, we performed a series post-alignment processing. First, mitochondrial sequences were excluded because no chromatin packaging exists [49]. We then sorted and converted files to .bam format using Samtools (version 1.11; Figure 3) [50]. The low-quality reads and redundancies generated during the PCR library building process were removed using Picard (version 2.25.4, https://broadinstitute.github.io/picard/). In order to ensure the correctness and rationality of biological conclusions, additional quality indicators need to be evaluated, such as mean insert size, corresponding standard deviation, TSS scanning score and fraction of reads in peaks (FRiP) score. The mean insertion size and corresponding standard deviation of paired-end were calculated with the Picard tool. TSS scanning score and FRiP were calculated with our own codes. Based on the quality control distribution of the population sample, we established the quality control characteristic threshold to filter a few samples with poor quality.

Finally, accessible chromatin regions were identified using MACS2 (version 2.2.7.1; Figure 3) [51, 52] based on peak calling with the parameter ‘--nomodel –shift -100 --extsize 200 -g 1.2e8’ [53-55]. The ‘-g’ parameter is effective genome size and was uniquely adjusted for each species. The example presents the ‘-g’ setting for *Arabidopsis thaliana*. These steps identified 18,065,954 ACRs from 638 samples. To assess the reliability of our settings, we compared the results with published data. Overall, we found that these ACRs were highly overlapped with the validated ACRs (overlap rate from 0.77 to 0.99, Supplement Table 2), suggesting that the peak calling for each species using the MACS2 tool is reliable. Additionally, we collected 273,472 validated ACRs from published reports [34, 56].

### ACRs visualization

To make it easier and more intuitive for users to compare different experimental data, we used the CHIPseeker toolset [57] to visualize ACRs for each sample, including the panorama distribution histogram of genome-wide ACRs, pie chart and combined charts of genome annotation, and heat map of ACRs near TSS. covplot function in CHIPseeker was used to visualize the panorama of ACRs distribution, which enables users to clearly observe the location of ACRs in the whole genome. AnnotatePeak function was used to analyze genome annotation, which classifies ACRs into promoter, 5’UTR, 3’UTR, 1st exon, other exon, 1st intron, other intron, downstream and distal intergenic. plotAnnoPie function was used to map genome annotation obtained by AnnotatePeak function into a pie chart. We set the 1kb upstream and downstream of TSS as window area, and used tagHeatmap function to draw a heat map of the ACRs combined with window area. Users can thus intuitively understand the distribution of ACRs near all gene promoters. Because ACRs location may not be unique (a ACR covers exons of one gene and, at the same time, introns of another gene), we drew the combined graph of venn pie and upset plot using UpSetR and vennpie functions.

### SNP annotation

In order to enable users to understand the relationship between chromatin accessibility and SNPs, we also downloaded SNPs of several existing species from Ensembl and Phytozome databases [58]. To achieve a high level of confidence in the variant calls from sequencing data, we applied a set of initial quality filters using VCFtools (version 0.1.16) [59] to filter out low quality SNPs. Specifically, we only kept variants with a mean read depth ≥ 10 to minimize spurious SNP calls due to low coverage genomic regions and with minor allele frequency (MAF) above 0.05 [60].

### Analysis of associated genes in chromatin accessibility regions

To further understand the function and transcriptional regulation of identified ACRs, PlantCADB classified associated genes for each ACR into three categories, i.e., Overlap genes, Closest genes, and Proximal genes. Overlap genes are defined as the genes that overlapped with the ACRs. The Closest genes are the ones closest to the center of each ACR. Proximal genes are defined as the genes in the upstream and downstream 1kb area of TSS that overlap with each ACR. The associated genes were downloaded from the Ensembl Plant and NCBI Genome databases. The specific classification method followed the steps in the ROSE script (ROSE_geneMapper.py) [61, 62]. On the query interface of PlantCADB, users can search data sets and corresponding regions based on the gene ID they are interested in.

### Analysis of transcriptional factors in chromatin accessibility regions

Analysis of transcription factor binding sites (TFBSs) in chromatin accessible landscapes reflects both aggregate TF binding and the regulatory potential of a genetic locus. PlantCADB used two methods to identify TFBSs: motif scan analysis and footprint analysis. We used MEME-suite [63] of FIMO tool [64] to scan for a single match of the motif of each ACR in each sample [56]. Plant-related TF lists and TF binding motifs for each species were obtained from PlantTFDB [65]. FIMO generates an ordered list of motifs as output, each with an associated log-likelihood ratio score, p-value and matching sequence. The log-likelihood ratio scores of each motif at each sequence position were calculated and converted into p-values by the dynamic programming. Users can set different thresholds to obtain the motif sequence.

Another way to decipher TF regulation rules is to use footprints. The combination of active TF and DNA will prevent enzyme from cutting at binding site, which is characteristic of ATAC-seq and DNase-seq experiments. This leads to objective formation of a protected area called footprint [66]. According to the characteristics of the two sequencing technologies and current footprint analysis tools, we used different analysis software for TF footprint analysis [49]. For ATAC-seq data, there are several obstacles in footprint analysis. First, due to the 9bp gap in the library construction process, displacement processing is required during the data handling process. In addition, Tn5 enzyme binding is biased, and the transient binding signal of TF is relatively weak. Among the existing ATAC-seq footprint analysis software, only HINT-ATAC can correct the cleavage preference of chain-specific Tn5 enzyme [67]. For the DNase-seq data, we used the HINT-BC tool to solve DNase-seq cleavage deviation and residence time that affect the calculation of footprint [68-70]. Both tools are based on hidden Markov model to predict TFBS with footprints, which outperform other tools.

After selecting appropriate methods, we downloaded position weight matrix (PWM) of motif from PlantTFDB database and created the regulatory genomics toolbox (RGT) data folder for each species. The folder had five files: gene, chromosome sizes, gene regions, annotation information and gene_alias. The gene_alias file allows for translation between multiple different gene IDs. The PWM was matched with the reference genome of the corresponding species, and resulted in protection score, tag count, number of binding sites and footprint logo combined with TF. Protection score measures the difference between enzyme digestion region counts in the flanking region and within the motif-predicted binding site. It has ability to detect TF with potential short residence time [69]. Tag count is used to represent the number of reads near the TFBSs that are ranked by footprint prediction. We also offer threshold conditions so that users can filter data according to their own criteria.

### Database and Web site implementation

The current version of PlantCADB is developed using Java 8 and HTML 5 and deployed to run on a Linux-based Apache Web server. The website page framework was designed and constructed by Bootstrap (Version 3.3.7), and the front and back data interaction was realized by JQuery (Version 3.6.0). Echarts (Version 3.7.0) was used to achieve data visualization. JBrowse2 browser framework was used for Genomic visualization. We recommend modern web browsers that support the HTML5 standard for the best display.

## Data availability

PlantCADB is freely available to the research community without login at https://bioinfor.nefu.edu.cn/PlantCADB/.

## CRediT author statement

**Ke Ding**: Methodology, Formal analysis, Data Curation, Conceptualization, Writing - original draft, Writing - review & editing. **Shanwen Sun**: Project administration, Writing - review & editing. **Chaoyue Long**: Software, Visualization. **Yang Luo**: Software, Visualization. **Jingwen Zhai**: Software. **Yixiao Zhai**: Visualization. **Guohua Wang**: Conceptualization, Supervision, Funding acquisition, Resources.

## Acknowledgements

This work was supported by The Innovation Project of State Key Laboratory of Tree Genetics and Breeding (Northeast Forestry University, No.2019A04), National Natural Science Foundation of China (No. 62001088, 62072095) and the Fundamental Research Funds for the Central Universities (No. 2572022BD04).

## Supplement figure legends

**Supplement figure 1 The relationship between ACRs and genome size**

**A**. The average number of ACRs per species (species are ordered based on the reference-genome size). **B**. Total sequence length of ACRs per species. **C**. The proportion of genome space with ACRs.

**Supplement figure 2 The relationship between ACRs types and gene density** The average percentage of dACRs, gACRs and pACRs in the genome of each species (species are ordered based on the gene density).

## Supplement table

**Supplement table 1 Statistics of reference genome Supplement table 2 Summary of PlantCADB’s data validation**

