## Supplementary figures and images for "PlantCADB: A comprehensive plant chromatin accessibility database"

### Supplement Figure 1

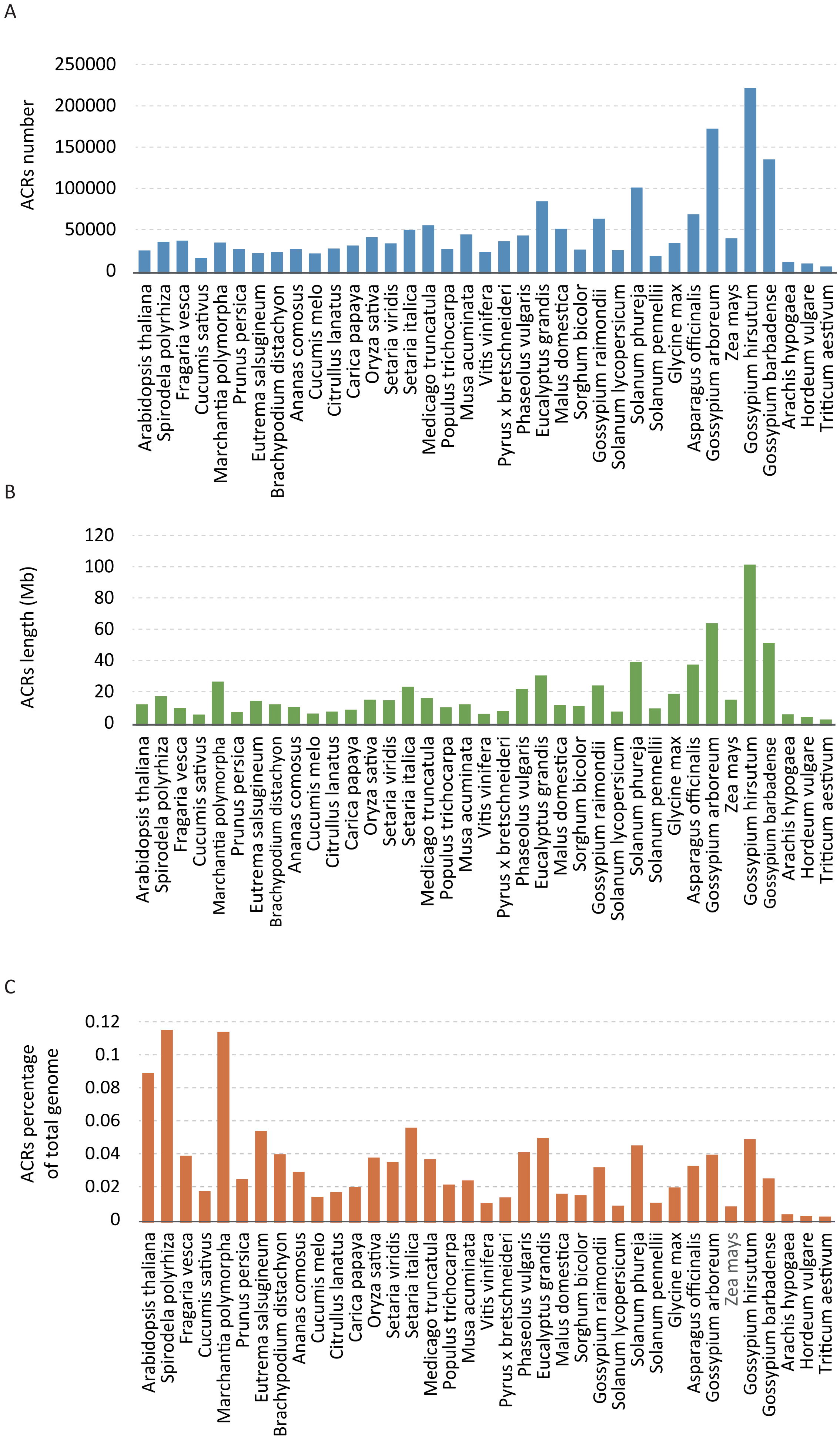


**Supplement Figure 1: The relationship between ACRs and genome size.**

### Supplement Figure 2

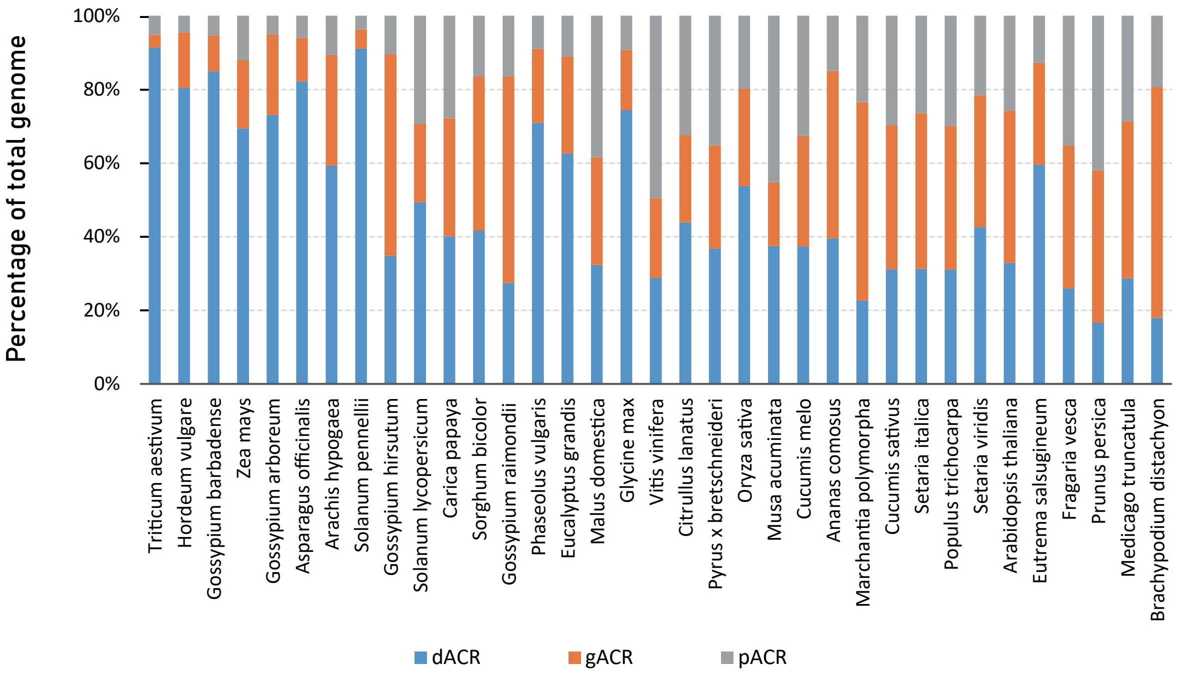


**Supplement Figure 2: The relationship between ACRs types and gene density.**
