## Supplement Table 1 for "PlantCADB: A comprehensive plant chromatin accessibility database"

**Supplement Table 1: Statistics of reference genome**

| **Species** | **Reference Genome** | **Size (byte)** | **Database** |
| --- | --- | --- | --- |
| *Ananas comosus* | F153 | 321104707 | Ensemble Plant |
| *Arabidopsis thaliana* | TAIR10 | 121662600 | Ensemble Plant |
| *Arachis hypogaea* | arahy.Tifrunner.gnm1.KYV3 | 2589093653 | NCBI_Genome |
| *Asparagus officinalis* | Aspof.V1 | 1203850602 | NCBI_Genome |
| *Brachypodium distachyon* | Brachypodium_distachyon_v3.0 | 275683706 | Ensemble Plant |
| *Carica papaya* | Papaya1.0 | 377124292 | NCBI_Genome |
| *Citrullus lanatus* | Cla97_v1 | 371549894 | Ensemble Plant |
| *Cucumis melo* | Melonv4 | 363822582 | Ensemble Plant |
| *Cucumis sativus* | Cucumber_9930_V3 | 229485246 | NCBI_Genome |
| *Eucalyptus grandis* | ASM1654582v1 | 624241621 | NCBI_Genome |
| *Eutrema salsugineum* | Eutsalg1_0 | 247207748 | Ensemble Plant |
| *Fragaria vesca* | FraVesHawaii_1.0 | 217450166 | NCBI_Genome |
| *Glycine max* | Glycine_max_v2.0 | 994826710 | Ensemble Plant |
| *Gossypium arboreum* | Gossypium_arboreum_v1.0 | 1727045129 | NCBI_Genome |
| *Gossypium barbadense* | Gossypium_barbadense_v1.1 | 2223503970 | NCBI_Genome |
| *Gossypium hirsutum* | Gossypium_hirsutum_v2.1 | 2217727374 | NCBI_Genome |
| *Gossypium raimondii* | Graimondii2_0 | 774180544 | Ensemble Plant |
| *Hordeum vulgare* | Hv_IBSC_PGSB_v2 | 4915007273 | Ensemble Plant |
| *Malus domestica* | ASM211411v1 | 714747307 | Ensemble Plant |
| *Marchantia polymorpha* | Marchanta_polymorpha_v1 | 229774095 | Ensemble Plant |
| *Medicago truncatula* | MedtrA17_4.0 | 419859025 | Ensemble Plant |
| *Musa acuminata* | MA1 | 480843747 | Ensemble Plant |
| *Oryza sativa* | IRGSP-1.0 | 381304565 | Ensemble Plant |
| *Phaseolus vulgaris* | PhaVulg1_0 | 529817333 | Ensemble Plant |
| *Populus trichocarpa* | Pop_tri_v3 | 441472116 | Ensemble Plant |
| *Prunus persica* | Prunus_persica_NCBIv2 | 231218665 | Ensemble Plant |
| *Pyrus x bretschneideri* | Pbr_v1.0 | 515220029 | NCBI_Genome |
| *Setaria italica* | Setaria_italica_v2.0 | 412522470 | Ensemble Plant |
| *Setaria viridis* | Setaria_viridis_v2.0 | 402328067 | Ensemble Plant |
| *Solanum lycopersicum* | SL2.50 | 837601628 | Ensemble Plant |
| *Solanum pennellii* | SPENNV200 | 938007464 | NCBI_Genome |
| *Solanum phureja* | ASM984975v1 | 900895428 | NCBI_Genome |
| *Sorghum bicolor* | NCBIv3 | 720622828 | Ensemble Plant |
| *Spirodela polyrhiza* | Spirodela polyrhiza 9509 v3 | 140326527 | NCBI_Genome |
| *Triticum aestivum* | IWGSC | 14789717189 | Ensemble Plant |
| *Vitis vinifera* | IGGP_12x | 494371940 | Ensemble Plant |
| *Zea mays* | AGPv4 | 2170684968 | Ensemble Plant |
