## Supplement Table 2 for "PlantCADB: A comprehensive plant chromatin accessibility database"

**Supplement Table 2: Summary of PlantCADB’s data validation**

| **Species** | **Sample ID** | **Ratio** | **Datasets** |
| --- | --- | --- | --- |
| *Arabidopsis thaliana* | Sample_00_057-Sample_00_059 | 0.99 | GSE128434[1] |
| *Brachypodium distachyon* | Sample_01_363-Sample_01_365 | 0.77 | GSE97369[2] |
| *Glycine max* | Sample_00_098-Sample_00_101 | 0.77 | GSE136645 |
| *Oryza sativa* | Sample_00_124-Sample_00_126 | 0.81 | GSE128434[1] |
| *Zea mays* | Sample_01_383-Sample_01_384 | 0.94 | GSE94291[3] |
|  | Sample_01_385-Sample_01_388 | 0.96 |  |
